# Reactivation of neural patterns during memory reinstatement supports encoding specificity

**DOI:** 10.1101/255166

**Authors:** Tobias Staudigl, Simon Hanslmayr

## Abstract

Encoding specificity or transfer appropriate processing state that memory benefits when items are encoded and retrieved in the same modality compared to when encoding and retrieval is conducted in different modalities. In neural terms, these effects can be expressed by a resonance process between a memory cue and a stored engram; the more the two overlap the better memory performance. We here used temporal pattern analysis in MEG to tap into this resonance process. We predicted that reactivation of sensory patterns established during encoding has opposing effects depending on whether there is a match or mismatch between the memory cue and the encoding modality. To test this prediction items were presented either visually or aurally during encoding and in a recognition test to create match (e.g. “dog” presented aurally during encoding and recognition) and mismatch conditions (e.g. “dog” presented aurally during encoding and shown visually during recognition). Memory performance was better for items in the match compared to the mismatch condition. MEG results showed that memory benefitted from neural pattern reinstatement only in the match condition, but suffered from reinstatement in the mismatch condition. These behavioural and neural effects were asymmetric in that they were only obtained for aurally encoded words but not for visually encoded words. A simple computational model was generated in order to simulate these opposing effects of neural pattern reactivation on memory performance. We argue that these results suggest that reactivation of neural patterns established during encoding underlies encoding specificity or transfer appropriate processing.

## Introduction

A memory cue is more effective when it overlaps with what has been studied. For instance, we are doing better in remembering when the context between encoding and retrieval is the same, compared to when they differ. A classic demonstration of this effect comes from Godden and Baddeley (Godden & Baddeley, 1975) who showed that the chances of retrieving a memory are higher when the environmental context during retrieval is the same as the one in which the item has been studied; i.e. items that are studied and retrieved on land (as opposed to when they are studied under water and retrieved on land). Similar effects have been observed with matching/non-matching background movies (Smith & Manzano, 2010; Staudigl, Vollmar, Noachtar, & Hanslmayr, 2015), or with different encoding/retrieval operations (i.e. rhyme vs semantic processing, (Morris, Bransford, & Franks, 1977). Similar effects have also been found with manipulating the modality in which items are presented at encoding and retrieval, for instance using pictures and words (Bauch & Otten, 2012; Mcdermott & Roediger, 1994), or presenting words visually and aurally (Mulligan & Osborn, 2009). In cognitive psychology these match/mismatch effects can be subsumed under the terms Encoding Specificity Principle (Tulving & Thomson, 1973), or Transfer Appropriate Processing (Morris et al., 1977). Both concepts highlight a basic organizing principle of human memory: The likelihood of remembering a certain memory increases with the degree to which a reminder resonates with a stored engram of that memory, i.e. interacts with a neural pattern that was established during encoding (Rugg, Johnson, Park, & Uncapher, 2008; Tulving, 1983).

Supporting evidence for a role of neural pattern reinstatement for memory retrieval comes from a number of studies showing reinstatement of encoding patterns during retrieval (Jafarpour, Fuentemilla, Horner, Penny, & Duzel, 2014; Johnson, McDuff, Rugg, & Norman, 2009; Michelmann, Bowman, & Hanslmayr, 2016; Polyn, Natu, Cohen, & Norman, 2005; Staresina, Henson, Kriegeskorte, & Alink, 2012; Staresina et al., 2016; Yaffe et al., 2014). These neural patterns can be detected using various multivariate analysis approaches in order to identify reactivation of neural patterns in space (Jafarpour et al., 2014; Polyn et al., 2005; Staresina et al., 2012), time (Michelmann et al., 2016; Staudigl et al., 2015), and time-frequency (Staresina et al., 2016; Yaffe et al., 2014). These studies demonstrate the general importance of neural pattern reinstatement for memory retrieval, however, we know little about the functional relevance of neural pattern reinstatement for memory and its involvement in Encoding Specificity (or Transfer Appropriate Processing). This is because there are hardly any studies which have tested the impact of neural pattern reinstatement on memory retrieval in the face of retrieval cues which match or do not match the encoding modality.

Two predictions can be made as to how neural pattern reinstatement impacts on memory in such matching and non-matching retrieval contexts. The first prediction is that the reinstatement of the original encoding pattern during retrieval is always beneficial for memory; regardless of a match between the retrieval and study contexts. In such a scenario, reactivation of the neural pattern containing the study episode does not support Encoding Specificity. A contrasting second prediction is that the reinstatement of encoding patterns is only beneficial for retrieval when the study and retrieval contexts match; if the study and retrieval context do not match, then reinstatement of encoding patterns might even be detrimental. In this second scenario, reactivation of the neural pattern containing the study episode would support Encoding Specificity. To the best of our knowledge, only one such study has been conducted which investigated neural pattern reinstatement in a paradigm using background movies as matching/non-matching contexts (Staudigl et al., 2015). Indeed, in this prior study we confirmed the second prediction, showing that memory benefitted from neural pattern reinstatement only when the encoding and retrieval contexts matched but suffered from reinstatement when the contexts did not match. This study can be seen as a first evidence that neural patterns established during encoding resonate with a retrieval cue, and that this resonance has a functional relevance for memory retrieval. This result therefore serves as a neural explanation for the Encoding Specificity Principle (or Transfer Appropriate Processing). However, if this is a general mechanism underlying Encoding Specificity, then we should be able to expand these results to different match/mismatch manipulations. Therefore, we here investigate neural pattern reactivation in a memory paradigm using a sensory modality match manipulation (Mulligan & Osborn, 2009) in which the items are studied and retrieved either visually or aurally (Figure 1A & Fig. 2).

**Figure 1.**
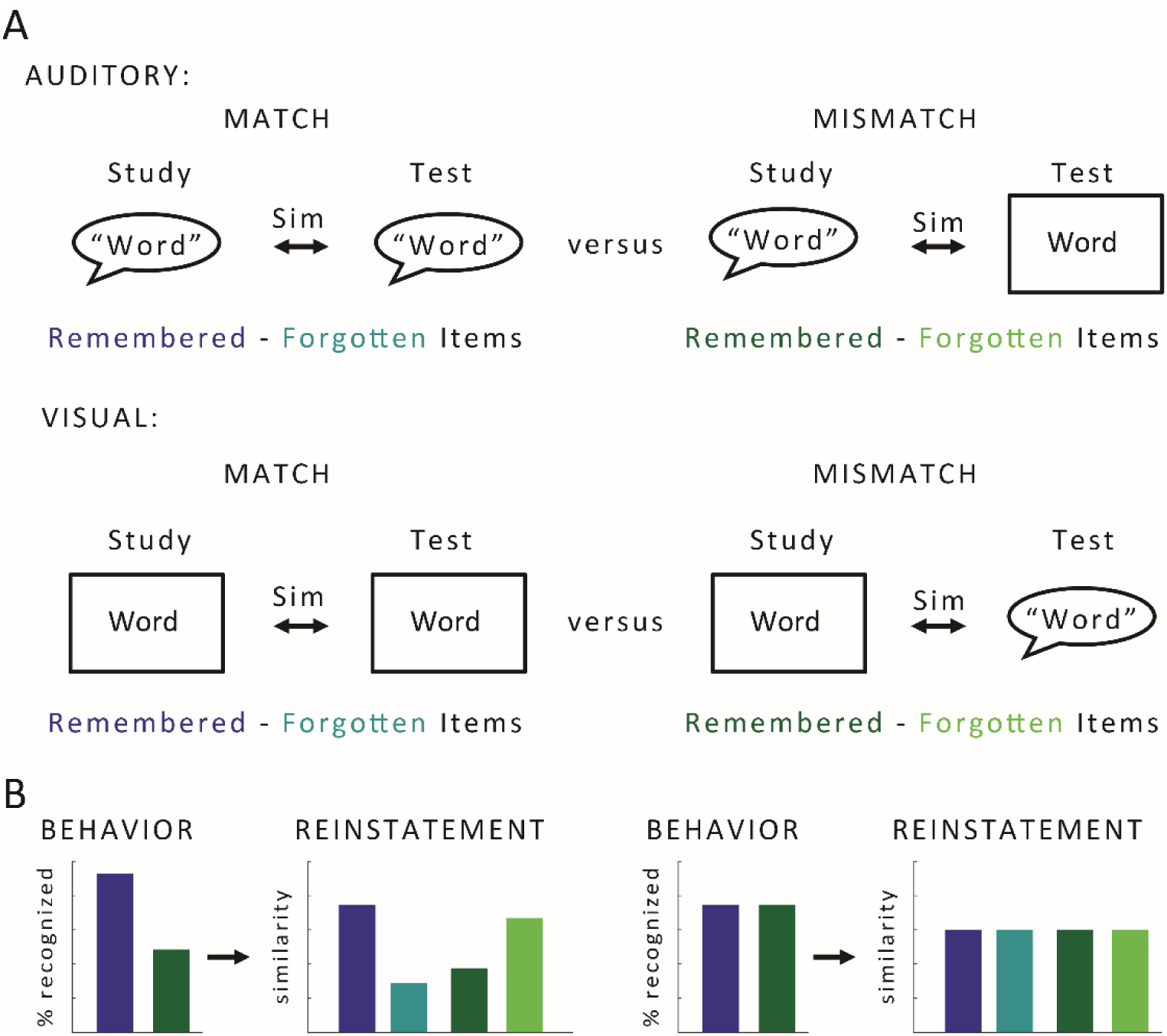
Analysis rationale and hypotheses. A) Temporal Pattern Similarity was computed for word pairs presented during study and test. Similarity differences between remembered and forgotten words were compared between the match and mismatch condition, separately for the auditory and the visual condition. B) A convergence between the behavioural results and the temporal pattern similarity is depicted. Left, if behaviour shows a match-mismatch effect we also expected an interaction effect between the differences (remembered – forgotten) in the match and mismatch condition (colour coded). Right, if no behavioural effect was observed then we also did not expect an interaction effect for temporal pattern similarity.

**Figure 2.**
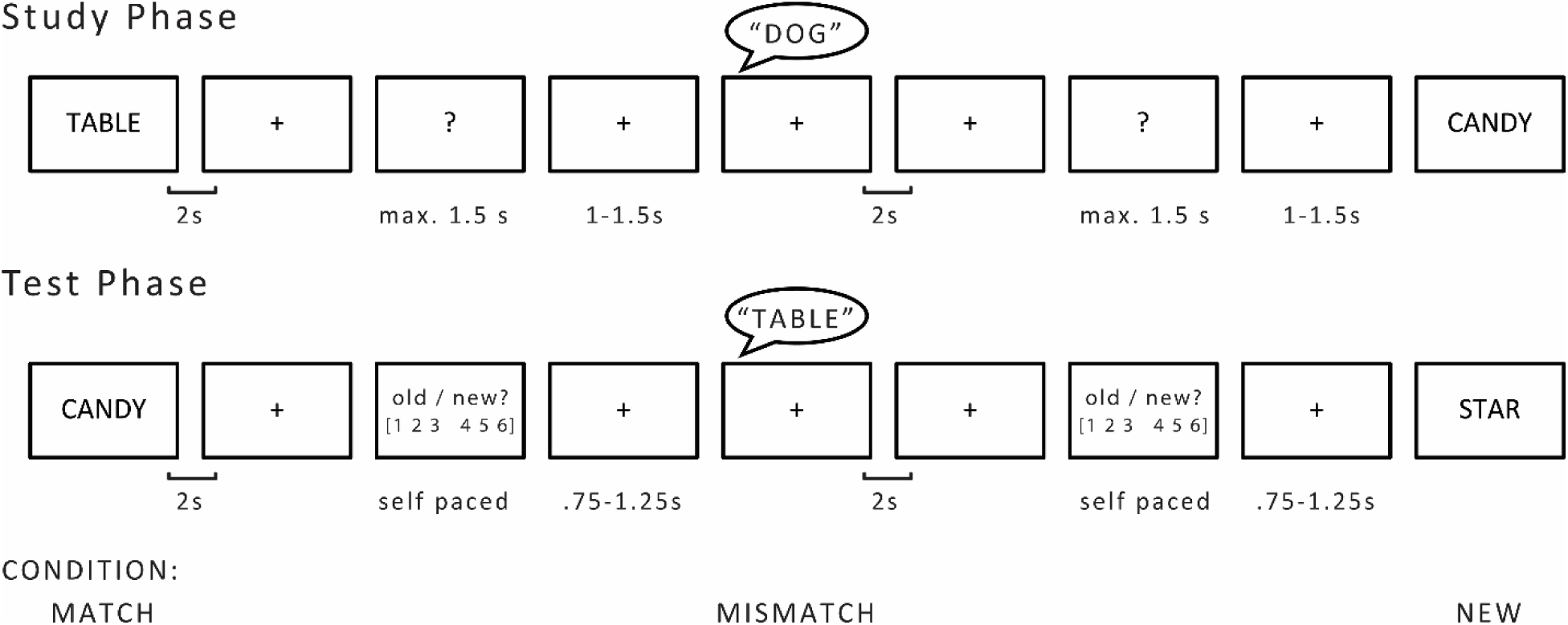
Experimental Procedure. In the study phase, words were either presented visually or aurally (indicated by the speech bubble). Duration of presentation was adjusted individually for each word to match the duration of the audio file. The combined presentation time of the word and fixation cross was 2 always seconds. Participants were instructed to count the syllables (two or more/less than two) of each word and respond during the question mark. In the test phase, old words were either presented in the same (match condition) or different (mismatch condition) modality as during the study phase. Old words were randomly intermixed with new words, and participants were asked to judge their confidence as to whether the word was old or new on a six-point scale, ranging from ‘very sure old’ (1) to ‘very sure new’ (6).

Neural pattern reinstatement can be measured in various ways with various recording techniques. Here we used MEG and focussed on the reinstatement of temporal patterns, as measured with phase in lower frequencies (<40 Hz). We chose this approach because previous studies showed that such phase based temporal similarity measures effectively captures the reactivation of temporal patterns in memory (Michelmann et al., 2016; Staudigl et al., 2015). We hypothesize to find higher temporal pattern similarity for successfully remembered items compared to forgotten items when the encoding and retrieval modalities match, and a reversal of this pattern when the encoding and retrieval modalities do not match. To foreshadow our findings, on a behavioural level we replicated the modality match effect (Mulligan & Osborn, 2009) for aurally encoded words, but not for visually encoded words (Fig. 3). Therefore, we analysed the MEG data separately based on encoding modality and expected the above described interaction pattern only for the condition which showed a behavioural effect (i.e. for aurally encoded words, Figure 1B).

**Figure 3.**
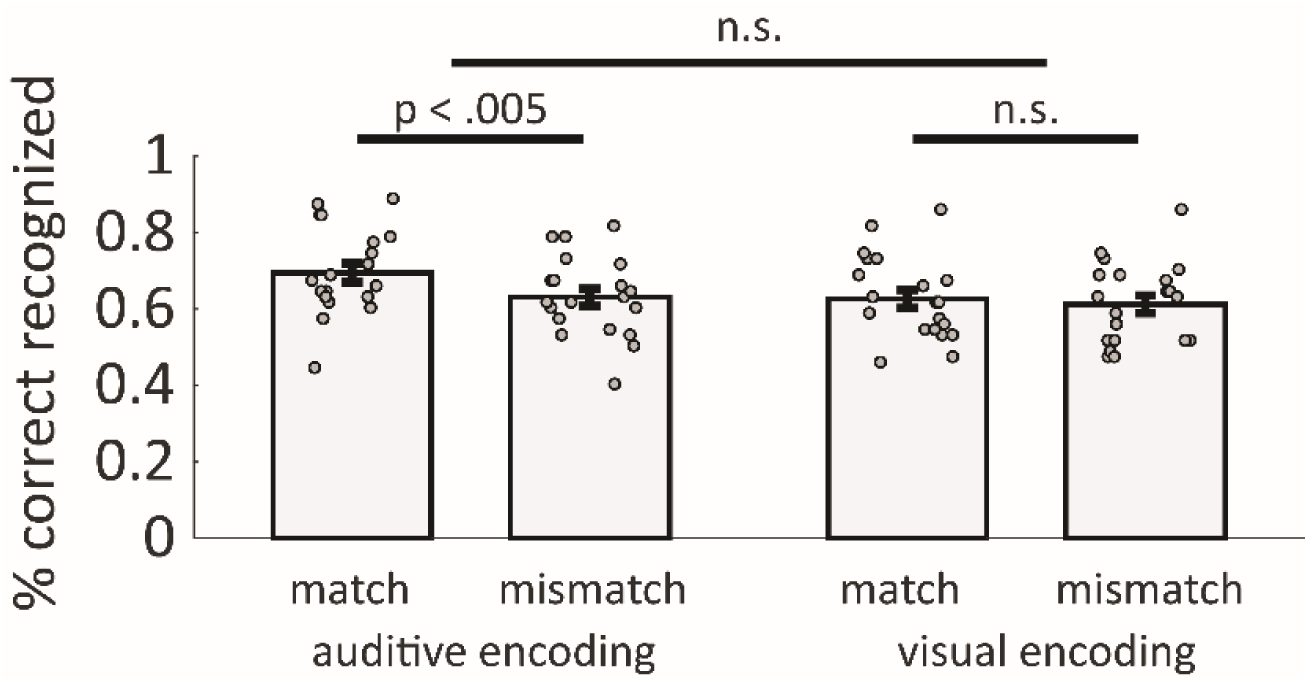
Behavioural results. Recognition performance in the match and mismatch condition as a function of auditory (left) and visual (right) encoding. A modality match effect was observed only for the auditory encoding condition (p < .005). Dots represent individual data points, errors bars depict S.E.

## Methods

### Ethics statement

The study was approved by the Institutional Review Board of the University of Konstanz. All participants gave written informed consent before the start of experiment in accordance with the Declaration of Helsinki.

### Participants

24 participants (between 19 and 26 years old; mean age = 22 years; 17 female; 21 right-handed) were recruited for study. After excluding four participants (technical problems, excessive environmental noise), data from 20 participants are presented here. Participants received course credits or monetary compensation for participation. All participants were German native speakers and reported no history of neurological disease and normal or corrected-to-normal vision.

Parts of this data have been published in (Staudigl & Hanslmayr, 2013) and (Westner, Dalal, Hanslmayr, & Staudigl, 2018), with respect to independent research questions and analyses.

### Procedure

The experiment consisted of a study and a test phase. An outline of the procedure is depicted in Figure 2. Before the start of the experiment, participants were instructed to count the syllables of each presented word and indicate via button press whether the word had two or more / less than two syllables. Participants were not instructed about the subsequent memory test, i.e., incidental encoding can be assumed. A short practice run ensured that the participants understood the task requirements.

During the study phase, words were presented in one of two modalities. In the visual encoding condition, words were projected on a screen. In the auditory encoding condition, vocal recordings of the words were presented via nonferromagnetic tubes to both ears, whilst the fixation cross remained on the screen. The duration of the word presentation was determined by the individual duration of the respective audio file, i.e., the time to pronounce the word (i.e. “dog” was presented for a shorter duration than “table”; mean duration = 697 ms, s.d. = 119 ms). Words were followed by a fixation cross. The duration of the word and fixation cross added up to 2000 ms. Thereafter, a question mark prompted the subject’s response in the syllable counting task. The presentation of the question mark was either ended by the subject’s button press or after a maximum duration of 1500 ms. A fixation cross (variable duration between 1000 and 1500 ms) preceded each word. After the study phase, participants performed a distracter task during which they counted backwards (in steps of three from a three-digit number) for 45 seconds. After the distracter task, participants were informed about the upcoming, surprise recognition memory test. Participants were instructed to indicate their confidence on whether the item was old (presented during the study phase) or new (not presented during the study phase) on a six-point scale ranging from ‘very sure old’ to ‘very sure new’. A short practice run ensured that the participants understood the task requirements.

During the test phase, words were presented with individual duration (determined by the duration of the respective audio file), followed by a fixation cross (summed duration of word and fixation cross = 2000 ms), followed by a stimulus picture depicting the response option. This stimulus prompted the participants’ response indicating their confidence on whether they recognized the word as old or new, by pressing one out of six specified buttons on the response panel. The stimulus picture prompting the response was shown until the participants gave a response. A fixation cross (variable duration between 750 and 1250 ms) preceded each word.

### Design and materials

420 unrelated German nouns were grouped into three lists with 140 words, each. Half of each list’s words had two syllables, the other half had one, three or four syllables. Two lists were presented during the study phase (old items). Half of these words were presented visually, the other half aurally. At test, all old items intermixed with the words from the remaining list (new items) were presented. Half of the old words were presented in the same modality (match condition), the other half in the different modality (mismatch condition) as during the study phase. Half of the new items were presented visually, the other half aurally.

The assignment of the lists to either study or test phase was counterbalanced across participants. For half of the participants, the confidence judgment during recognition test ranged from ‘very sure old’ (1) to ‘very sure new’ (6), for the other half from ‘very sure new’ (1) to ‘very sure old’ (6), thereby counterbalancing the response lateralization across subjects. Items were presented in a random order, with the constraint that not more than five words of the same modality (visual, auditory) and not more than five words from the same condition (match, mismatch) were presented sequentially.

### Data acquisition and preprocessing

MEG data was recorded with a whole-brain 148-channel magnetometer system (MAGNESTM 2500 WH, 4D Neuroimaging, San Diego, USA) inside a magnetically shielded room from participants in supine position. The data was continuously recorded at a sampling rate of 678.17 Hz and bandwidth of 0:1-200 Hz. Preprocessing of the data was done using the fieldtrip toolbox (Oostenveld, Fries, Maris, & Schoffelen, 2011), an open-source MATLAB toolbox for MEEG data analysis. Data from the study and test phase was epoched into single trials, with epochs ranging from 1500 ms prior to the onset of word presentation to 4000 ms after onset of word presentation. Trials were visually inspected for artefacts, to reject contaminated trials. Thereafter, independent component analysis (ICA) was used to correct for blinks, eye movements, and cardiac artefacts.

All trials remaining after artefact inspection were categorized according to the behavioural performance of each participant’s during the recognition test phase. Trials including old items that were confidently judged as old (responses 1, 2, and 3) constituted remembered trials, the remaining trials including old items were classified as forgotten trials, in both the study and the test phase. Trials including new items that were confidently judged as being new (responses 4, 5, and 6) constituted correct rejections, the remaining trials including new items were classified as false alarms.

### Similarity analysis

In order to identify reinstatement of neuronal patterns from encoding during retrieval, similarity in neuromagnetic activity between words presented at study and test was assessed. Based on previous findings (Michelmann et al., 2016; Staudigl et al., 2015), we focused on similarity in oscillatory phase, using the pairwise phase consistency index (PPC; (Vinck, van Wingerden, Womelsdorf, Fries, & Pennartz, 2010). PPC estimates the phase consistency between two separate signals across trials, quantifying the extent of a consistent phase relationship among them. Compared to other measures of phase consistency (e.g., phase-locking value), PPC is advantageous since it is not biased by the number of trials.

In order to provide the necessary phase information for the computation of the PPC, time-frequency representations of the data were computed by a sliding time window approach with a window length of 0.5 s in steps of 50 ms across the data. After multiplying a hanning taper to each window, the Fourier transformation was calculated for frequencies between 2 and 40 Hz in steps of 2 Hz.

The PPC was computed between the same words from the study and the test phase, separately for remembered and forgotten trials, match and mismatch condition, and auditory and visual encoding condition.

### Statistics

Statistical quantification of the data was performed by a cluster-based nonparametric permutation approach (Maris & Oostenveld, 2007) identifying clusters of activity on the basis of rejecting the null hypothesis while controlling for multiple comparisons over sensors, time-points and frequencies. For each time and frequency bin at each sensor, a test statistic was calculated (10,000 permutations), based on a paired samples t-test comparing the difference in PCC in the match condition (remembered minus forgotten trials) to the difference in PPC in the mismatch condition (remembered minus forgotten trials; see Figure 1).

T-values above the cluster-forming threshold (p < 0.05, two-sided t-test) were clustered based on adjacency in time, frequency and space (a minimum of 2 adjacent sensors was required for forming a cluster). T-statistics were summed in each cluster and compared against the distribution of maximal clusters provided by the permutation approach. Only the cluster with the largest summed value was considered and tested against the permutation distribution. The null-hypothesis that the match and mismatch condition showed no difference in PPC was rejected at an alpha-level of 0.05 (two-tailed).

### Simulation

A simple computational model was generated to illustrate the opposing effects that the reactivation of neural patterns may have on memory performance in match and mismatch conditions. The model was inspired by a theoretical paper by E. Tulving (Tulving, 1983) who envisioned an early resonance process, termed “Ecphory” which can be described mathematically via a correlation between a stored engram and a retrieval cue and which inputs into the retrieval process. The simulation was programmed in MATLAB (version R2016a) and is a slightly modified version of a previous simulation published in (Staudigl et al., 2015). The code for the simulation is available on https://github.com/hanslmayr/Audivis_Simulation. The model is divided into two layers, a semantic layer and a sensory layer, to simulate item meaning and the sensory modality in which an item is presented. Therefore, the sensory layer is further divided into auditory and visual units (see Figure 5). The semantic layer has 12 units; the sensory layer had 6 units for visual and 6 units for auditory patterns. Item patterns were expressed by a combination of 3 activated units on each layer (see Figure 5); 100 patterns were generated to simulate encoding and retrieval of 100 items. Thereby, each item was represented by a unique combination of a 3 unit pattern in a semantic layer and a 3 unit pattern in a sensory layer. These patterns were expressed in forms of 3 by 4 matrices, containing zeros for non-activated units and ones for activated units (see Figure 5).

Memory retrieval was simulated via an interaction, i.e. correlation, between a *reactivated pattern (R*) and a *cued pattern (C*) (Tulving, 1983). The simulation was calculated individually for each item, resulting in 100 trials. Within each trial, the reactivated pattern was first calculated separately for the semantic and the sensory layer. On the semantic layer, the reactivated semantic pattern (*R_sem_*) was simply assumed to be the same as the cued pattern (*C_sem_*) with a constant small amount of white noise (range 0 to 0.1). No difference was assumed between match and mismatch conditions, as in our experiment items were identical on a semantic level between match and mismatch conditions. On the sensory layer, we calculated the reactivated pattern (*R_sens_*) by taking the original sensory pattern and adding white noise onto it. The strength of noise was controlled by a reactivation parameter in order to simulate the effects of strength of reactivation on memory performance (i.e. high levels of noise result in low levels of reactivation). The cued pattern (*C_sens_*) on the sensory layer varied between match and mismatch conditions. For the match condition, *C_sens_* was the original pattern in the respective sensory layer in which the item has been encoded (i.e. auditory layer for aurally encoded words). Therefore, for match conditions *C_sens_* overlapped with *R_sens_*, with the strength of this overlap depending on the noise parameter which controls the strength of reactivation. For mismatch conditions, *C_sens_* was taken from a different units of the sensory layer (i.e. visual units for aurally encoded words). Therefore, for mismatch conditions *C_sens_* never overlapped with *R_sens_*. Finally, memory performance (*Mem*) for a given trial was calculated by averaging the 2d correlations (r) between the cued and reactivated patterns on the semantic and sensory layers as per the below equation.

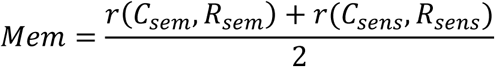

## Results

### Behavioural results

A summary of the behavioural results is depicted in Fig. 3. Averaged across modality conditions, participants correctly recognized significantly more words (t_19_ = 3.56, p < 0.005) in the match (mean = .661, SE = .023) than the mismatch condition (mean = .621, SE = .020). The interaction between modality (auditory vs. visual), test condition (match vs. mismatch) was not significant (F_1,19_ = 2.72, p > .115).

In the auditory encoding condition, participants correctly recognized significantly more words in the match (mean = .695, SE = .025) than in the mismatch (mean = .63, SE = .023) condition (t_19_ = 3.55, p < .005). No significant difference between correctly recognized words in the match (mean =.626, SE = .024) and the mismatch (mean = .612, SE = .024) condition was found for visually encoded items (t_19_ = .73, p > 0.475).

The false alarm rate was not significantly different (t_19_ = −0.62, p > .54) for items presented visually (mean = .26, SE = .027) or aurally (mean = .27, SE = .025) during test.

### Reinstatement of auditory temporal patterns interacts with modality match

In order to identify memory reinstatement of temporal patterns from encoding during retrieval, similarity in neuromagnetic activity between words presented at study and test was assessed using the pairwise phase similarity (Michelmann et al., 2016). Given the asymmetric behavioural pattern, i.e. a modality match effect was only observed for aurally encoded words, the analysis was carried out separately for visually and aurally encoded words.

In the auditory encoding condition, cluster-based nonparametric statistics yielded a significant interaction effect (p cluster < .05), indicating that the difference in the match condition (remembered – forgotten) was reliably higher than in the mismatch condition (remembered – forgotten) at 6-8 Hz between 0.15 and .2 seconds after word onset (Fig. 4a). Figure 4b depicts a topography of the interaction effect, highlighting left central sensors contributing to the significant cluster.

**Figure 4.**
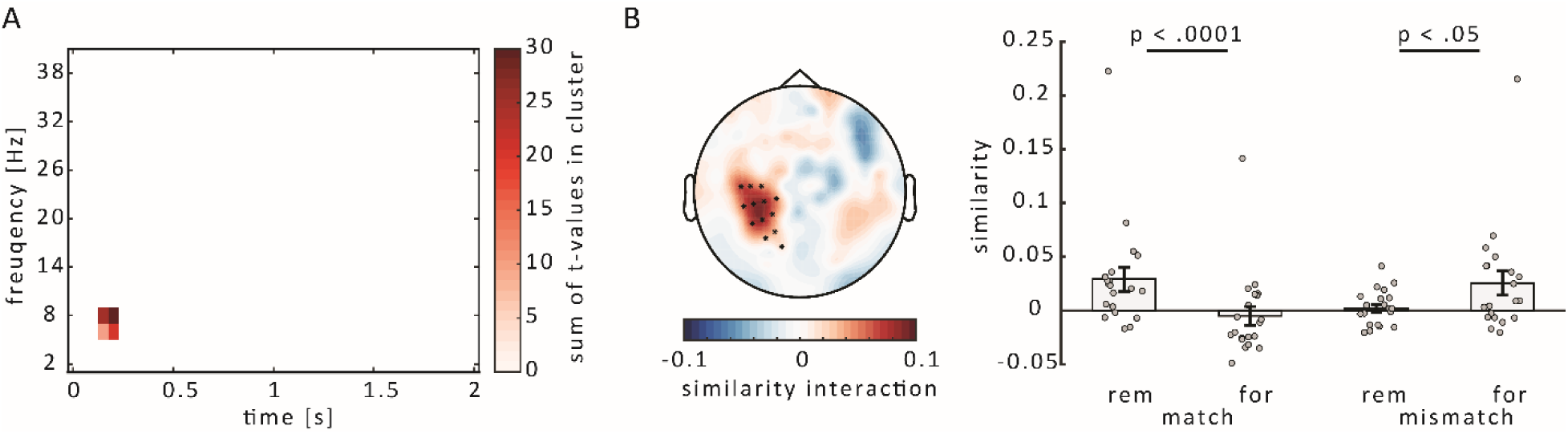
Interaction effect in the auditory condition. A) Significant interaction (p cluster < .05) between the match (remembered – forgotten) and the mismatch (remembered – forgotten) condition, at 6-8 Hz and .15-.2 s after word onset. B) Left: Topography of the interaction effect, averaged across time and frequency depicted in A. Sensors contributing to the significant interaction are highlighted. Right: Similarity for the remembered and forgotten words, in the match and mismatch condition, respectively, as averaged across time (.15-.2 s), frequencies (6-8 Hz) and sensors highlighted in the topography. Dots represent individual data points, errors bars depict S.E.

Post-hoc t-tests revealed that the neuronal similarity was higher for the remembered than the forgotten words in the match condition for (t_19_ = 5.02, p < .0001), whereas in the mismatch condition, similarity was higher for the forgotten than the remembered words (t_19_ = 2.11, p < .05, see Figure 4b). It should be noted that this post-hoc analysis is statistically biased by selectively averaging across those channels, time and frequency bins that show a significant interaction.

Nevertheless, this post-hoc analysis adds information as it shows that the interaction is not just driven by a difference in one condition, but by opposing differences between remembered and forgotten items in both conditions.

In the visual condition, there were no significant interaction effects when comparing the difference (remembered – forgotten) in the match condition to the difference (remembered – forgotten) in the mismatch condition (all cluster p’s > .59).

When remembered versus forgotten items were compared irrespective of match / mismatch condition, no significant effects were found in the auditory (p cluster > .08) nor in the visual condition (p cluster > .16).

### Simulating the effects of high vs low reactivation on memory performance

The results above confirm our hypothesis that the reactivation of encoding patterns is beneficial or detrimental for memory retrieval depending on whether the retrieval cue matches the modality of encoding or not. We propose that these results are suggestive of a resonance process between a retrieval cue and the reactivated encoding pattern (i.e. the engram) which determines the memory outcome (Tulving, 1983). In order to illustrate this process we constructed a simple computational model (see Methods), which is illustrated in Figure 5. This model is a modified version of a model we’ve published in a previous study (Staudigl et al., 2015). Items were generated in terms of patterns in two different layers, a semantic layer, which represents the meaning of the item, and a sensory layer, representing the sensory features of the item. The sensory layer contains visual and auditory units. Memory performance is conceived of as a correlation between the input pattern, represented by the cue, and the reactivated encoding pattern. This correlation between an input pattern and a stored engram describes the resonance, ecphory-like, process between a cue and an item stored in memory (Tulving, 1983). The stronger the resonance, the more likely an item will be endorsed as “old”. In the simulation we varied the level of reactivation (i.e. high vs low) of the encoding pattern on the sensory layer only, since item meaning was kept constant in our experiment. The results of this simple model capture the empirical findings obtained for the auditory encoding condition. Specifically, if the modality of the retrieval cue matches the encoding modality, high levels of sensory reactivation lead to a strong resonance, i.e. a better memory performance. On the other hand, if the modality of the retrieval cue does not match the modality of the encoded pattern, then high levels of sensory reactivation lead to lower resonance, i.e. worse memory performance. This latter effect arises because in the mismatch modality the patterns of the engram and the cued patterns never overlap as they are represented in different units. Reactivation of a sensory trace in the mismatch condition therefore leads to a negative correlation which then reduces the overall correlation between the cue and the reactivated pattern.

**Figure 5.**
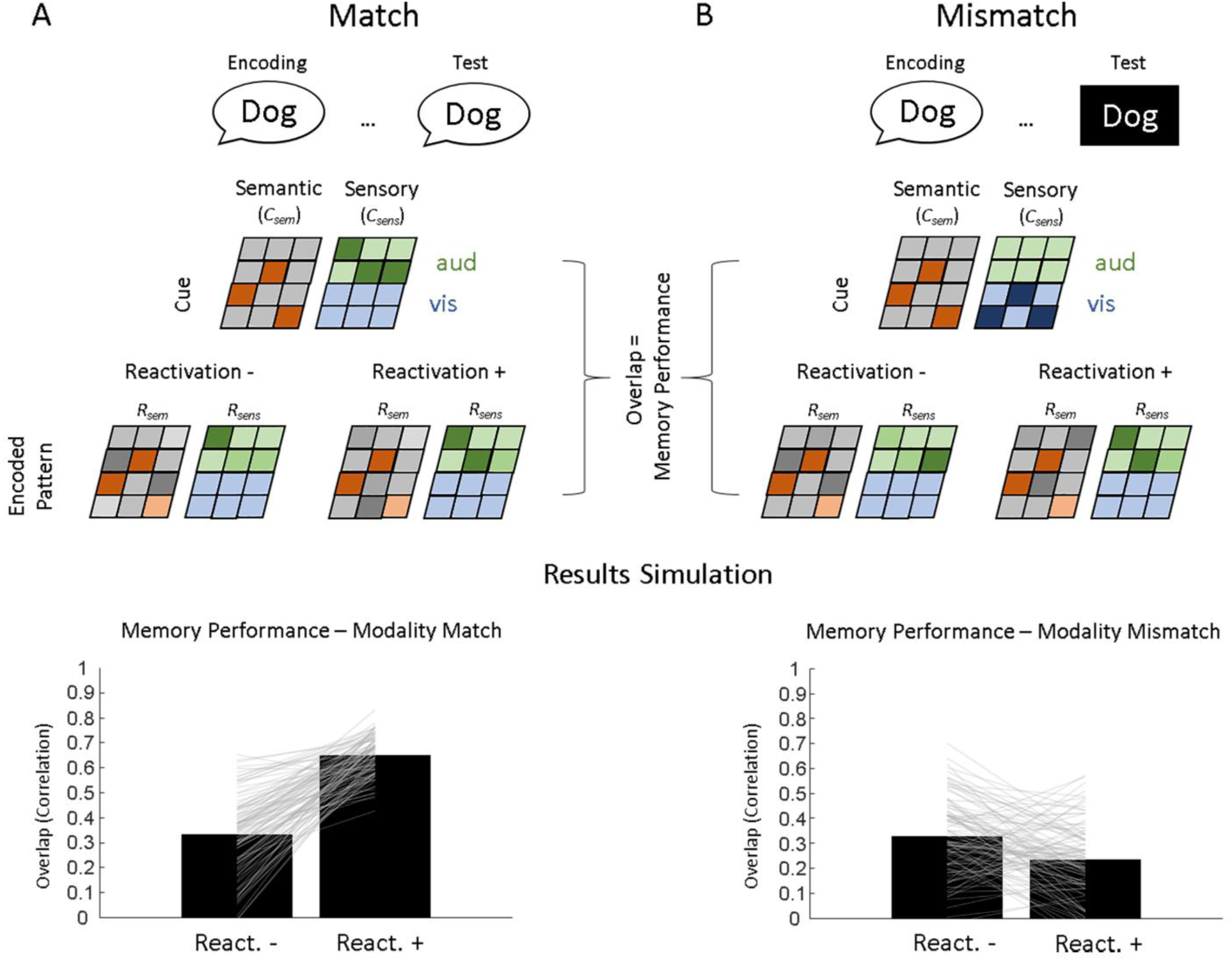
The architecture of a simple computational model (top panel) and results of the simulation (bottom panel) are shown for Match (A) and Mismatch (B) conditions. Individual patterns were generated on a semantic and a sensory layer to represent cued and encoded item patterns. The sensory layer was divided into visual and auditory units. Memory performance was conceived of as the overlap between a cued pattern (*C_sem_*, *C_sens_*) and a reactivated pattern (*R_sem_, R_sens_*) as calculated with 2d correlations. These correlations were averaged across the two layers. The bottom panel shows the overlap averaged across simulation runs (bar plots) and individual trials (grey lines) for high and low reactivation levels (React.+ and React.-, respectively). 100 simulation runs per condition and reactivation level were carried out.

## Discussion

In the current study we investigated how the reactivation of encoding patterns interacts with memory in the face of retrieval cues which match or do not match the encoding modality. For aurally encoded words, we show that reactivation of encoding patterns is related to opposing memory outcomes; depending on whether the retrieval cue matches the encoding modality or not. Specifically, reactivation of a sensory encoding pattern is beneficial for memory when the retrieval cue matches the encoding modality, but is detrimental when the cue does not match the encoding modality. Together with our previous findings (Staudigl et al., 2015), these results provide insights into the neural basis underlying a classic effect in cognitive psychology, i.e. Encoding Specificity or Transfer Appropriate Processing (Morris et al., 1977; Tulving & Thomson, 1973): the more the retrieval cue overlaps with the encoded pattern, i.e. the stronger the two resonate, the more likely it is that the item will be recognized as old (Rugg et al., 2008). Paradoxically, when the retrieval cue is presented in a different sensory modality than in which the item has been encoded, reactivation of sensory patterns was related to decreased memory performance. This pattern can be explained on the basis of a simple model which illustrates our results on a neural and behavioural level: the mismatch between the reactivated sensory pattern (i.e. “dog” in auditory layer) and the sensory pattern provided by the retrieval cue (i.e. “dog” visually presented) decreases the overlap (or resonance) between the cue and the reactivated pattern.

Neural evidence consistent with Encoding Specificity comes from fMRI (Park & Rugg, 2008) and ERP studies (Bauch & Otten, 2012). These studies show that the neural correlates of successful encoding vary depending on whether the memory is later tested with cues that match or do not match the encoding modality (Park & Rugg, 2008). We here add to this neural evidence by using a temporal pattern similarity approach and show that reactivation of neural patterns established at encoding has opposing effects on memory depending on how the memory is cued. One advantage of the here used approach is that it returns a time-resolved measure of memory reactivation, which is allows to infer whether reactivation is an early process leading up to a memory judgement or whether it is a later by-product of a memory judgement (i.e. imagery, or retrieval monitoring). The early time window of reactivation observed here (i.e. 150 – 200 ms) supports the former view and suggests an early resonance process between a memory cue and a stored engram which occurs before the memory decision is being made, which has been termed Ecphory (Tulving, 1983). This finding is consistent with a larger body of studies describing similar early reactivation effects (Jafarpour et al., 2014; Waldhauser, Braun, & Hanslmayr, 2016; Wimber, Maass, Staudigl, Richardson-Klavehn, & Hanslmayr, 2012).

Interestingly, on a behavioural level the observed modality match effects were asymmetric, i.e. a modality match effect was observed only for aurally encoded words but not for visually encoded words (albeit it should be acknowledged that there was no significant 2-way interaction between match/mismatch and modality). Consistent with this asymmetric pattern, only aurally encoded items showed an interaction effect in terms of neural pattern reactivation with modality match, whereas no such effect was obtained for visually encoded items. A possible post-hoc explanation for this asymmetry is that the visual presentation of words automatically evokes auditory patterns encoded by the participants. This would be consistent with reports of high levels of auditory cortex activation typically observed during reading, and in line with the idea that auditory cortex is part of the brain’s “reading network” (Wandell & Le, 2017). This activation of auditory patterns might have been further promoted by the here used encoding strategy, i.e. syllable counting which arguably enforces subjects to covertly pronounce the word. In such a scenario, for a visually encoded word, an auditory cue is just as effective as a visual cue because the engram contains both patterns. Consistent with this explanation, a previous study (Mulligan & Osborn, 2009) which reported modality match effects for visually *and* aurally encoded words used a different encoding strategy, i.e. intentional encoding (as opposed to syllable counting and incidental encoding employed in our study). On a more general level, these considerations highlight the importance of the cognitive processes carried out at encoding and retrieval which determine memory effects observed on a behavioural and neural level (Hanslmayr & Staudigl, 2014; Rugg et al., 2008).

The here observed reactivation of auditory patterns occurred in a frequency range of 6-8 Hz. This frequency range perfectly matches a previous study where we found that dynamic auditory patterns are encoded in the phase of a 6-8 Hz oscillation (Michelmann et al., 2016). This result is also in line with another study showing that auditory stimuli can be decoded from neural phase patterns at 4-8 Hz (Ng, Logothetis, & Kayser, 2013). This result is, however, in contrast to a previous study (Staudigl et al., 2015) investigating match-mismatch with background movies, where we found reactivated patterns at a higher frequency range (i.e. 30 Hz). This difference could be either due to the different sensory modalities of the stimuli between the two studies (i.e. auditory vs visual), or could be due to slightly different analysis approaches. In contrast to Staudigl et al. 2015, we here used pairwise phase consistency (PPC) as we did in Michelmann et al (2016) which has the advantage of resulting in a time-resolved measure of Temporal Pattern Similarity. Together with previous studies (Michelmann et al., 2016; Ng et al., 2013), and another study in this same special focus (Michelmann et al., submitted), the results suggest a general role of 6-8 Hz oscillations for coding auditory information not only on a perceptual level, but also in memory.

## Conclusion

The encoding specificity principle or transfer appropriate processing are classic frameworks which have influenced memory research for decades. In their seminal paper published in 1973, Tulving and Thomson describe the basic ideas of Encoding Specificity but also state: *„The terms are ill defined, and the concepts do not explain too much at this time. Yet they serve to remind us that something else besides the properties of a presented item determines how well the item is remembered and that an important research problem is to find you what this something else is and how it works” (Tulving & Thomson, 1973).*

More than 4 decades later we still know little about the mechanisms underlying the encoding specificity principle. In neural terms, the encoding specificity principle can be expressed as a resonance process between a cue and a stored engram, whereby the overlap between the two determines whether a memory can be retrieved or not (Rugg et al., 2008; Tulving, 1983). We here show that the strength of reactivation of that memory trace plays a central role in this process and thus is a critical ingredient underlying Encoding Specificity (or Transfer Appropriate Processing). We hope that future studies pick up on these ideas using multivariate analysis tools which allow to quantify the reactivation of stored memory traces (i.e. the engrams) using standard neuroimaging tools such as EEG, MEG or fMRI.

## Acknowledgements

We thank Ann-Kristin Rombach, Leona Hellwig, Marina Koepfer, and Janine Weichert for help with data acquisition. This research was supported by European Union’s Horizon 2020, (https://ec.europa.eu/programmes/horizon2020/, 661373) to TS; Deutsche Forschungsgemeinschaft (http://www.dfg.de/, Emmy Noether Programme Grant HA 5622/1-1) to SH; European Research Council (https://erc.europa.eu/, Consolidator Grant 647954) to SH; Wolfson Foundation and Royal Society (https://royalsociety.org/grants-schemes-awards/grants/wolfson-research-merit/) to SH.

## Author Contributions

T.S. and S.H. designed the experiment and wrote the paper. T.S. collected the data. T.S. performed the analyses, S.H. performed the simulation.

## Competing Financial Interest

The authors declare no competing financial interests.

